# Deficient Executive Control in Transformer Attention

**DOI:** 10.1101/2025.01.22.634394

**Authors:** Suketu Patel, Hongbin Wang, Jin Fan

**Affiliations:** Department of Psychology, Queens College, The City University of New York, Queens, NY 11367, USA; College of Medicine, Texas A&M University, Houston, TX 77030, USA

**Keywords:** Transformer, Large Language Model (LLM), Attention, Executive Control, Conflict Effect, Stroop Task

## Abstract

Although transformers in the large language models (LLMs) effectively implement a self-attention mechanism that has revolutionized natural language processing, they lack an explicit implementation of executive control of attention found in humans, which is essential for resolving conflicts and selecting relevant information in the presence of competing stimuli, and is critical for adaptive behavior. To investigate this limitation in LLMs, we employed the classic color Stroop task that is widely regarded as the gold standard for testing executive control of attention. Our results revealed a typical conflict effect of better performance in terms of accuracy in the congruent condition (e.g., naming the ink color of the word RED in red) compared to the incongruent condition (e.g., naming the ink color of the word RED in blue), which is similar to human performance in short sequences. However, as sequence length increased, the performance degraded toward chance levels on the incongruent trials despite maintaining excellent performance on congruent trials and near-perfect word reading ability. These findings demonstrate that while transformer attention mechanisms can achieve human-comparable performance in smaller contexts, they are fundamentally limited in their capacity for conflict resolution across extended contexts. This study suggests that incorporating executive control mechanisms akin to those in biological attention could be crucial for achieving more general reasoning and reliable performance toward artificial general intelligence.

## Introduction

The introduction of the Transformer model in their paper "Attention is All You Need" (Vaswani et al., 2017) marked a significant milestone in the recent transformative development of machine learning (ML) and artificial intelligence (AI). This model, which relies on an attention mechanism instead of recurrent or convolutional layers to uncover essential spatiotemporal structures in input data, has become the foundation for state-of-the-art natural language processing (NLP) models. Since then, the attention mechanism has been applied to various multimodal large language models (LLMs) in tasks beyond NLP, including computer vision, speech recognition, and even video generation (OpenAI, 2023; Anthropic, 2024). The revolutionary success of transformer-based LLMs naturally invites a critical question regarding the role of attention in ML and AI, including its nature, boundaries, and limitations. In particular, how is transformer-based machine attention different from human attention, which has been a fundamental concept in psychology and has been actively studied for millennia?

William James famously stated "Everyone knows what attention is. It is the taking possession by the mind, in clear and vivid form, of one out of what seem several simultaneously possible objects or trains of thought" (James, 1890). However, modern neuroscience regards attention not as a simple and single mechanism but as a complex mental faculty subserved by multiple neuronal networks. According to attention network theory (ANT), for example, there are three different attentional functions of alerting, orienting, and executive control, each based on a distinct neuronal network, that can be differentiated (Posner & Petersen, 1990; Fan, McCandliss, Sommer, Raz, & Posner, 2002). Could a single mechanism of transformer attention possibly encompass the whole spectrum of biological attention? Here we show that transformer attention primarily corresponds to the orienting function of biological human attention and falls short in realizing other types of attention. In particular, using a classic Stroop effect paradigm, we demonstrate that even the state-of-the-art LLMs (ChatGPT 4o and Claude 3.5) show deficient executive control performance. We discuss that such analysis and comparison are important for us to seek artificial general intelligence (AGI).

Human attention is complex, and many different models are introduced to explain the mechanisms. The ANT proposes that the three networks, alerting, orienting, and executive control, are functionally integrated and interact to selectively focus cognitive resources on specific stimuli, tasks, or information while filtering out irrelevant or less important inputs (Posner & Petersen, 1990). The alerting network is responsible for achieving and maintaining an alert state with brain areas such as the locus coeruleus involved. The orienting network is involved in selecting specific information from the sensory input, which includes directing attention to a particular stimulus or location. Key areas involved in orienting include the superior colliculus, frontal eye fields, and parietal lobe. An executive control network is engaged in tasks that require control over thoughts and actions in the presence of competing computations of information and is essential to adaptive behavior. Regions of the frontoparietal network, including the anterior cingulate cortex (ACC), the anterior insular cortex, and the dorsolateral prefrontal cortex (DLPFC), are involved (Fan et al., 2003; Petersen & Posner, 2012).

In addition to its explanatory relevance in biological attention, the ANT has been reliability-validated neural functionally within healthy aging clinical research (Ishigami et al., 2016), pathologies, and attention deficit hyperactivity (Gradys, Lipowska, Bieleninik, & Dzhambov, 2021). The ANT has also been computationally validated through connectionist modeling (Wang & Fan, 2007). Behavior and functional neuroimaging studies have demonstrated that these three attentional networks function independently and interactively (Fan, McCandliss, Sommer, Raz, & Posner, 2002; Fan, McCandliss, Fossella, Flombaum, & Posner, 2005; Fan, Gu, Guise, Liu, Fossella, Wang, & Posner, 2009; Xuan et al., 2016). Selective impairment of the executive function may result in mental disorders with deficient conflict processing (Wang et al., 2005).

Modern neural network-based AI systems can efficiently process large amounts of input information distributed spatially (e.g., in image processing) or temporally (e.g., in NLP) to reach decisions, such as image classification or next-word prediction. The essence of the attention mechanism, as implemented in transformers, lies in learning a function that facilitates the extraction and selection of the most relevant information from the input, allowing the model to "attend" to it. Briefly, a transformer model segments the input into a sequence of tokens, encoding each token as an embedded vector in a high-dimensional vector space. To predict the next token given an input sequence (i.e., a prompt), the model employs a learned attention function to assign weights to each input token and make predictions based on their weighted sum. Compared to traditional ML techniques, a key distinguishing feature of the transformer attention mechanism is its ability to prioritize relevant input tokens by assigning weights based on latent semantic information rather than relying solely on simple spatial or temporal proximity.

In modern multimodal LLMs, which can process input across different modalities (e.g., visual, audio, text, video), tokens from different modalities are represented and cross-referenced using a unified embedding space (e.g., Radford et al., 2021; Dosovitskiy et al., 2021; OpenAI, 2023). In multimodal systems, visual inputs typically undergo preprocessing through vision transformers (ViT) or convolutional neural networks to create patch embeddings that can be processed alongside text tokens (Dosovitskiy et al., 2021). These visual embeddings are then projected into a shared latent space where they can interact with text embeddings through self-attention mechanisms (Li et al., 2022). The cross-modal attention architecture typically employs a multi-head attention mechanism where some heads might focus on object-word associations while others attend to spatial relationships or abstract concepts (Khan et al., 2023). This parallel processing allows the model to capture complex relationships between modalities, similar to how human perception integrates multiple sensory inputs (Stein, Stanford, & Rowland, 2020). It is clear that in both canonical and multimodal transformers, the attention mechanism primarily corresponds to the orienting function of human attention in the ANT framework as long as we expand the scope of orienting to include cross-modal semantic dimensions beyond spatiotemporal factors. However, is this type of attention all we need? Is it sufficient to account for other ingredients of human attention functions required for natural intelligence as suggested by the ANT?

While these systems implement sophisticated forms of self-attention and cross-modal attention, the executive control component analogous to the anterior cingulate and lateral prefrontal cortices has not been explicitly implemented. Experiments that test the conflict effect could be conducted to verify this claim by showing results indicating whether transformer-based attention produces human or superhuman performance or if there are deficits compared to the biological attention. Standard psychological tasks could also help to quantify these differences, particularly in measuring how transformers handle attentional conflicts. The Stroop effect, first documented by John Ridley Stroop in 1935, is a robust psychological phenomenon that demonstrates interference in the reaction time and accuracy of a task when there is a mismatch between the physical color of a word and the meaning of the word (Stroop, 1935; MacLeod, 1991). This interference effect (or called conflict effect) occurs when individuals must name the ink color of a word while ignoring the word’s meaning. For example, when participants are asked to name the ink color of the word "BLUE" printed in red ink, the incongruent condition, they typically respond more slowly and make more errors compared to when the word and ink color match, e.g., “RED”, the congruent condition. This effect highlights the automatic nature of reading and the cognitive effort required to override this automatic process when naming colors, providing insights into attention, processing speed, and executive function (Cohen, Dunbar, & McClelland, 1990). Within biological attention, the Stroop effect has been utilized to test the conflict effect in the human brain (e.g., Leung et al., 2000; Fan, Flombaum, McCandliss, Thomas, & Posner, 2003). Because the Stroop effect has systematic variation across conditions and persists even with practicing the tasks, human performance suggests it reflects fundamental aspects of executive control of attention.

In this study, we investigated the Stroop effect in two state-of-the-art LLMs with multimodal transformers: OpenAI’s GPT-4o and Anthropic’s Claude 3.5 Sonnet. To assess the models’ conflict resolution capabilities, we presented lists of words as visual images to the models under different conditions (congruent, incongruent, and neutral control) and asked them to report either the ink color names or the words themselves. A key experimental manipulation involved progressively increasing the word list length from 1 to 40 items. Model performance was evaluated based on accuracy (rather than reaction time, given the online nature of these experiments). Would the models exhibit a conflict effect similar to that observed in humans? We predicted that the models would demonstrate the Stroop effect and that the models would exhibit behavior patterns distinct from those of humans. Collectively, these findings would highlight the limitations of LLMs in executive control of attention.

## Method

Two state-of-the-art (SOTA) multi-modal LLMs, ChatGPT (Generative Pre-trained Transformer) 4o and Claude Sonnet 3.5, were evaluated for their executive control of attention capabilities using a modified Stroop task paradigm. ChatGPT has the architecture of a decoder-only transformer with everything related to the encoder removed and is pre-trained via self-supervised learning over large amounts of text data. Claude Sonnet 3.5, like other LLMs, is built on a transformer architecture. It also relies on decoder-only transformer architectures rather than incorporating an encoder and uses self-attention mechanisms in the decoder to process input sequences and generate outputs, enabling efficient text generation and completion tasks.

Transformers are particularly well-suited for NLP due to their ability to handle long-range dependencies in text through mechanisms like self-attention and positional encoding.

### Stimulus materials

The stimuli presented to the AI models for the color naming task and the word reading task were the following five conditions (Figure 1, also see Supplementary Table 1):

(1) Congruent condition: Color words printed in their corresponding colors, e.g., RED.
(2) Incongruent condition: Color words printed in non-matching colors, e.g., RED.
(3) Mixed condition: 50% of congruent and 50% incongruent color-word trials.
(4) Neutral condition: Common office-related words matched for character length with color words, e.g., PEN.
(5) Nonword neutral condition: To test the potential interference of word meaning on color naming, an additional nonword neutral control condition with the character of Xs (e.g., XXX) matching the length of the color words was included only for the color naming task.

**Figure 1.**
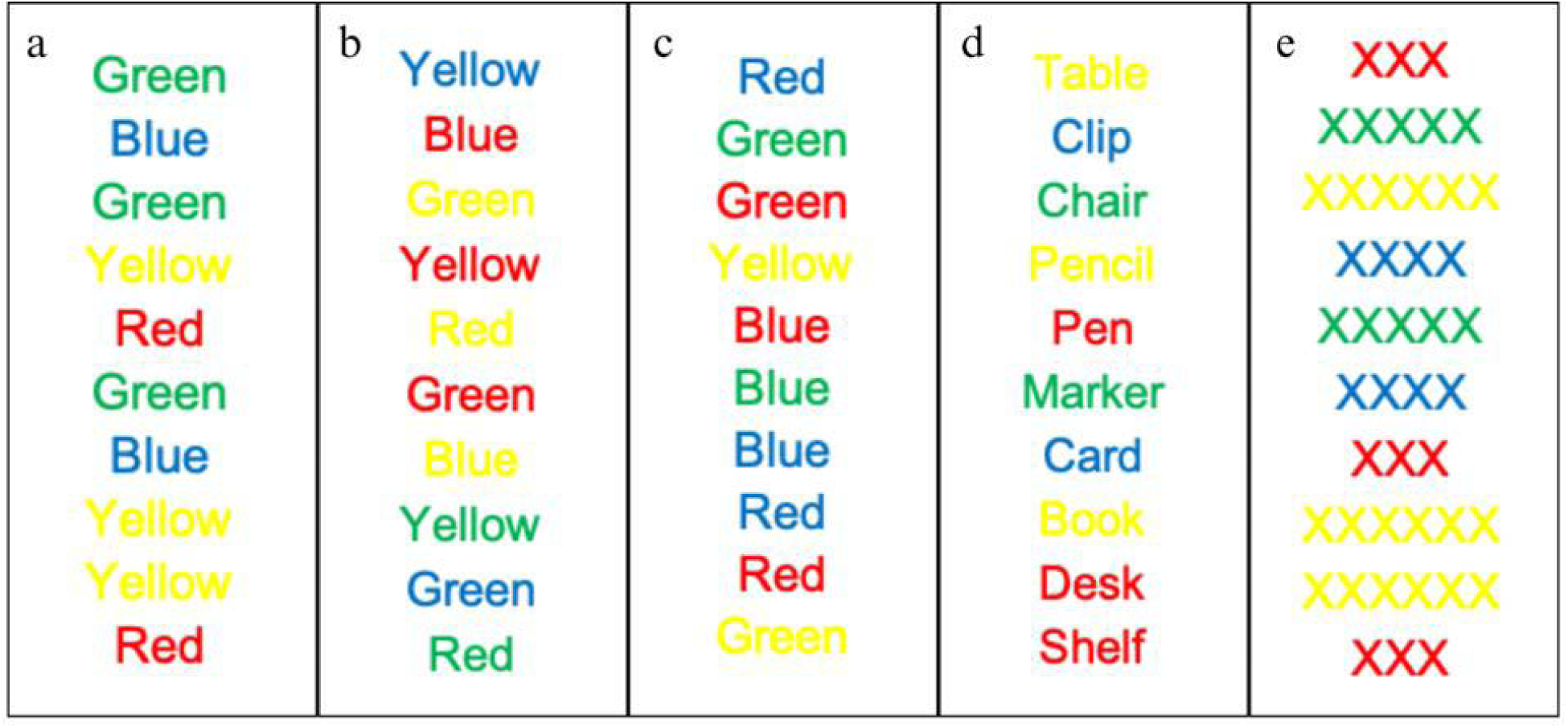
Illustration of the color word lists of congruent (a), incongruent (b), mixed (c), and neutral (d) conditions for the length of 10 words. An additional control condition, the nonword neutral condition is also illustrated (e).

The stimuli were composed of the following four hex colors and words: RED (#EA3323), YELLOW (#FFFF54), GREEN (#4EAE5B), and BLUE (#0070C0). For the neutral condition, the following words were used: BOOK, DESK, CARD, PEN, TABLE, PENCIL, SHELF, MARKER, CLIP, CHAIR. The stimuli text used Arial font and was composed of vertically arranged word lists created with Microsoft Excel. They were randomized using a random number generator to uniquely re-order each stimuli list (see Experimental Tasks below).

### Experimental tasks

The study comprised two tasks of color naming and word reading. The first task is a standard color naming Stroop task utilizing a within-subjects design with the five conditions. The second task is a standard word reading task, which also utilizes a within-subjects design with the four conditions without the nonword neutral condition. The same stimuli were used in the two tasks with the 5 list lengths: 1, 5, 10, 20, and 40 words. The only variation in the word reading task was the prompt used alongside the word list (see below).

### The Stroop effect

The Stroop effect also called the conflict effect or interference effect, is a widely used measure in cognitive psychology and neuropsychology. This effect is defined as the performance (e.g., in terms of accuracy or reaction time) difference between the incongruent and congruent conditions, or between the incongruent and neutral conditions, in the color naming task. For this study, the conflict effect was also examined in the mixed condition for the accuracy difference between incongruent and congruent trials.

### Procedure

All stimuli were presented as PNG image files containing vertically arranged word lists. Each Stroop task trial was initialized in a unique chat instance, where the prompt was provided with an attached image of the list of words. For the five list lengths: 1, 5, 10, 20, and 40 words, each list length had 30 trials and was repeated for both LLMs. For each of the 30 test trials for the five list lengths, a single chat session was utilized to test each of the conditions. The output was then compared to the original list, and manual scoring and verification were conducted to count how many answers were correct. This Stroop paradigm with the two tasks was repeated for each of the LLMs tested. Both LLMs were presented with identical stimulus images across conditions, and a new session was initiated for each set of four conditions to prevent context carryover. The task order of the conditions was varied in each trial to counter order effects.

The following multi-modal (Google, 2024) prompt was used for the color naming task: “For each word presented in this image, respond with ONLY the color the word is written in. Do not explain your reasoning or add any other words. Respond as quickly as possible with just the color.”

For the word reading task, the instruction was: “For each word presented in this image, respond with ONLY the word that is written. Do not explain your reasoning or add any other words. Respond as quickly as possible with just the word.”

### Data analysis

Response coding followed a systematic protocol where one-to-one mapping was employed when output word count matched input word count, with responses mapped sequentially. In cases where output exceeded input, responses were paired based on semantic correspondence to the label and color. For incomplete responses where outputs contained fewer words than inputs, optimal matching was employed to maximize accuracy assessment.

Responses were coded as incorrect if they contained both the color name and word label, and sequential matching was employed with each output response paired with the next word in the input sequence. Missing responses were coded as incorrect responses and skipped so that the LLMs were not penalized for responses after the omission.

Getting the exact number of words in the output as specified in the input is non-trivia, and often, for example, in a 20-word list, there would be outputs that ranged from 15-25 words. By chunking groups of correct answers and skipping over repeated, best extrapolations are used to decide if responses were missed or if they are incorrect.

The trials were analyzed for average correct number, percent correct (accuracy), error rate, standard deviation (STD), and standard error (SE). For the performance of the LLMs, human performance patterns were compared or referenced (see Supplementary Table 1).

## Results

### Color naming task results

The observed performance patterns broadly parallel those found in human results (MacLeod, 1991). Like humans, both LLMs demonstrated relatively high performance (in terms of accuracy) for the word reading task and reduced performance in the incongruent condition compared to congruent and neutral conditions for the color naming task (Tables 1&2 and Figures 2&3). Within the two task formats, computerized item-by-item or blocked on paper, only older adults had degraded performance on the blocked version, while younger adults maintained performance across both versions (Ludwig et al., 2010). This parallel is particularly relevant for evaluating LLMs, which undergo blocked testing in list lengths of 5, 10, 20, and 40 words, as well as item-by-item testing in single word trials. Unlike humans, though, who can maintain a a 95% accuracy and a consistent reaction time across trials that span 5, 10, 20 (Dallaway, Lucas, & Ring, 2023) and 97% accuracy in 60-minute color reading trials with 1500 word list length (Salihu et al., 2023) (see Supplementary Table 1), LLMs’ performance generally declined as a function of the length of the word list length. There was one notable exception in Claude 3.5 Sonnet’s congruent condition between 10 and 20 words, where accuracy improved from 90% to 99%. This anomaly can be attributed to several trials in the 10-word condition yielding zero correct answers, thereby lowering the mean performance.

**Table 1.** Results of color naming task (Mean, SD, and SE)

| Condition | Word List Length | Percent Correct - GPT 4o | Percent Correct - Sonnet 3.5 | ChatGPT 4o | Claude Sonnet 3.5 |
| --- | --- | --- | --- | --- | --- |
| Congruent | 1 | 100% | 83% | 1 (0) [0] | 0.83 (0.38) [0.07] |
| Congruent | 5 | 100% | 100% | 5 (0) [0] | 5 (0) [0] |
| Congruent | 10 | 99% | 90% | 9.93 (0.25) [0.05] | 9 (3.05) [0.56] |
| Congruent | 20 | 99% | 99% | 19.83 (0.59) [0.11] | 19.73 (1.28) [0.23] |
| Congruent | 40 | 89% | 92% | 35.73 (5.69) [1.04] | 36.87 (4.58) [0.84] |
| Incongruent | 1 | 100% | 100% | 1 (0) [0] | 1 (0) [0] |
| Incongruent | 5 | 91% | 97% | 4.57 (0.57) [0.1] | 4.87 (0.35) [0.06] |
| Incongruent | 10 | 57% | 75% | 5.7 (1.7) [0.31] | 7.5 (2.03) [0.37] |
| Incongruent | 20 | 22% | 76% | 4.4 (2.61) [0.48] | 15.13 (3.42) [0.62] |
| Incongruent | 40 | 15% | 24% | 6.17 (3.85) [0.7] | 9.4 (4.95) [0.9] |
| Mix | 1 | 100% | 97% | 1 (0) [0] | 0.97 (0.18) [0.03] |
| Mix | 5 | 99% | 99% | 4.93 (0.25) [0.05] | 4.93 (0.25) [0.05] |
| Mix | 10 | 72% | 79% | 7.23 (1.72) [0.31] | 7.93 (1.78) [0.32] |
| Mix | 20 | 52% | 78% | 10.47 (1.31) [0.24] | 15.63 (2.57) [0.47] |
| Mix | 40 | 41% | 50% | 16.4 (4.55) [0.83] | 19.9 (4.08) [0.74] |
| Neutral | 1 | 100% | 73% | 1 (0) [0] | 0.73 (0.45) [0.08] |
| Neutral | 5 | 99% | 100% | 4.97 (0.18) [0.03] | 5 (0) [0] |
| Neutral | 10 | 94% | 96% | 9.37 (1.43) [0.26] | 9.63 (0.76) [0.14] |
| Neutral | 20 | 75% | 78% | 14.9 (2.66) [0.49] | 15.63 (2.57) [0.47] |
| Neutral | 40 | 32% | 27% | 12.73 (4.71) [0.86] | 10.87 (3.17) [0.58] |
| Nonword Neutral | 1 | 100% | 100% | 1 (0) [0] | 1 (0) [0] |
| Nonword Neutral | 5 | 97% | 100% | 4.83 (0.46) [0.08] | 5 (0) [0] |
| Nonword Neutral | 10 | 85% | 100% | 8.53 (1.14) [0.21] | 10 (0) [0] |
| Nonword Neutral | 20 | 76% | 100% | 15.17 (1.72) [0.31] | 20 (0) [0] |
| Nonword Neutral | 40 | 58% | 100% | 23.07 (3.43) [0.63] | 40 (0) [0] |

**Figure 2.**
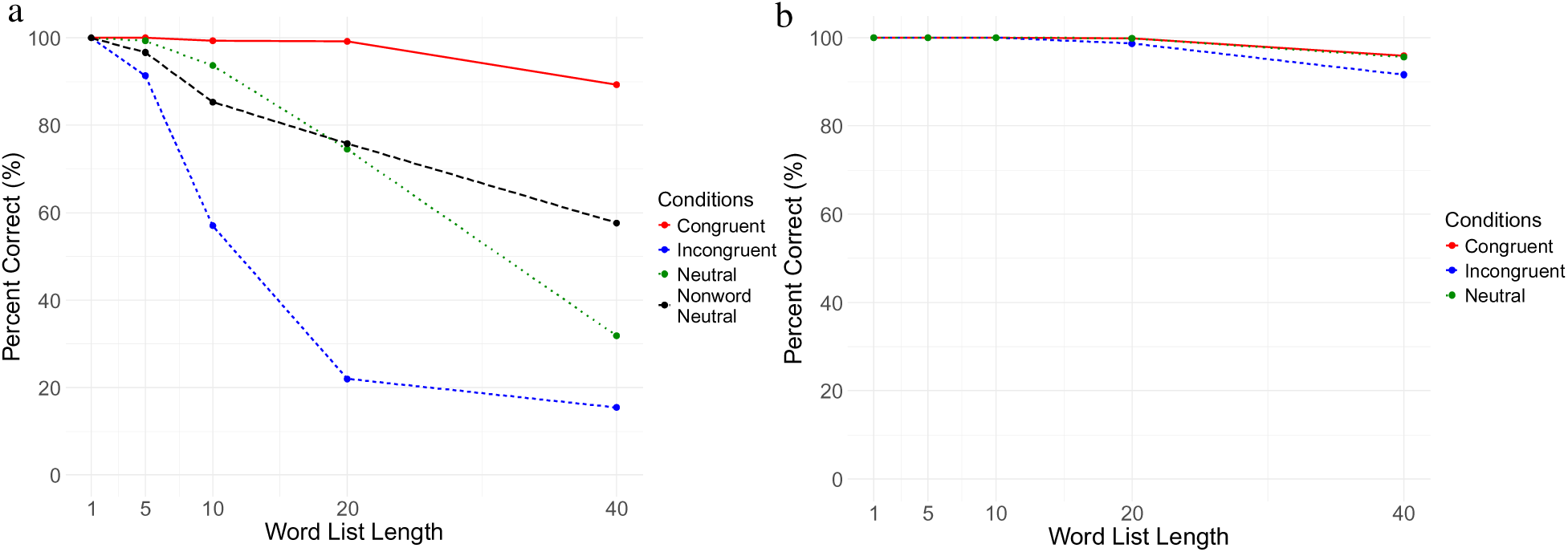
Percent correct for all conditions by word list length (ChatGPT 4o). (a) color naming; (b): word reading.

For the color naming task, the most substantial performance degradation occurred between 20 and 40 words across most conditions except in the congruent condition. For the congruent condition, both models maintained robust performance, with GPT-4o and Claude 3.5 Sonnet achieving 99% accuracy at 20 words, declining modestly to 89% and 92%, respectively, at 40 words. The neutral condition exhibited more pronounced degradation, with both models showing approximately a 20% decrease from 10 to 20 words, followed by a 50% reduction from 20 to 40 words.

The incongruent condition revealed unique model-specific differences in degradation patterns. GPT-4o demonstrated early performance deterioration, with accuracy dropping from 91% at 5 words to 57% at 10 words, followed by further substantial declines to 22% at 20 words and to 15% at 40 words (Figure 2a). In marked contrast, Claude 3.5 Sonnet maintained relatively stable performance through 20 words (76% accuracy) before experiencing a sharp decline to 24% accuracy at 40 words, a 52% drop in performance (Figure 3a). The divergent pattern suggests some functional differences in how the two models handle increased cognitive load under incongruent conditions. The significant degradation pattern of the two LLMs suggests fundamental limitations compared to biological attention.

**Figure 3.**
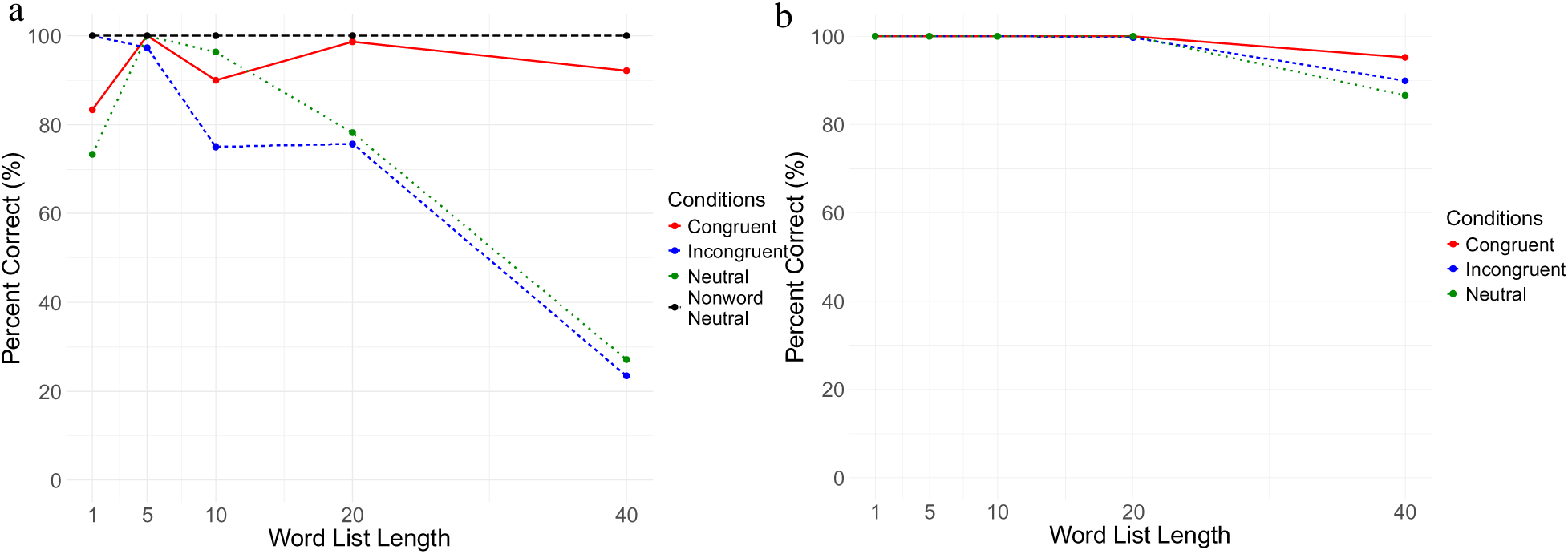
Percent correct for all conditions by word list length (Claude Sonnet 3.5). (a) color naming; (b) word reading.

In the mixed trial condition (Table 3, Figures 4&5), where the list consisted of an equal split between congruent and incongruent trials (50% each) but in random order, GPT-4o exhibited above-chance performance only at shorter list lengths (100% at 1 word, 99% at 5 words, and 72% at 10 words), approaching chance levels of 50% (assuming if each response is solely based on word meaning) at 20 words (52%) and falling below chance at 40 words (15%) for color naming (Figure 4a). The separation of the two trial types showed a predictable pattern of accuracy distribution between congruent and incongruent trials. Performance was nearly perfect or high accuracy for congruent trials (e.g., 100% for 1 word list and 81% for 40 word list), while errors and extremely low accuracy were exclusively observed in incongruent conditions for longer lists (i.e., 10, 20, and 40). Specifically, GPT-4o’s performance on incongruent trials showed a marked decline as list length increased, reduced to 46% accuracy with 10-word lists and dropping dramatically to just 1% accuracy for both 20- and 40-word lists. Remarkably, in the incongruent condition, performance deteriorated to nearly 0%, indicating a complete breakdown of executive control, a pattern that stands in stark contrast to typical human performance, where individuals maintain significant, though reduced, high accuracy despite an interference effect (see Supplementary Table 1). This near-total performance collapse suggests a fundamental limitation in Chat GPT4o’s capacity for conflict resolution and response inhibition.

**Figure 4.**
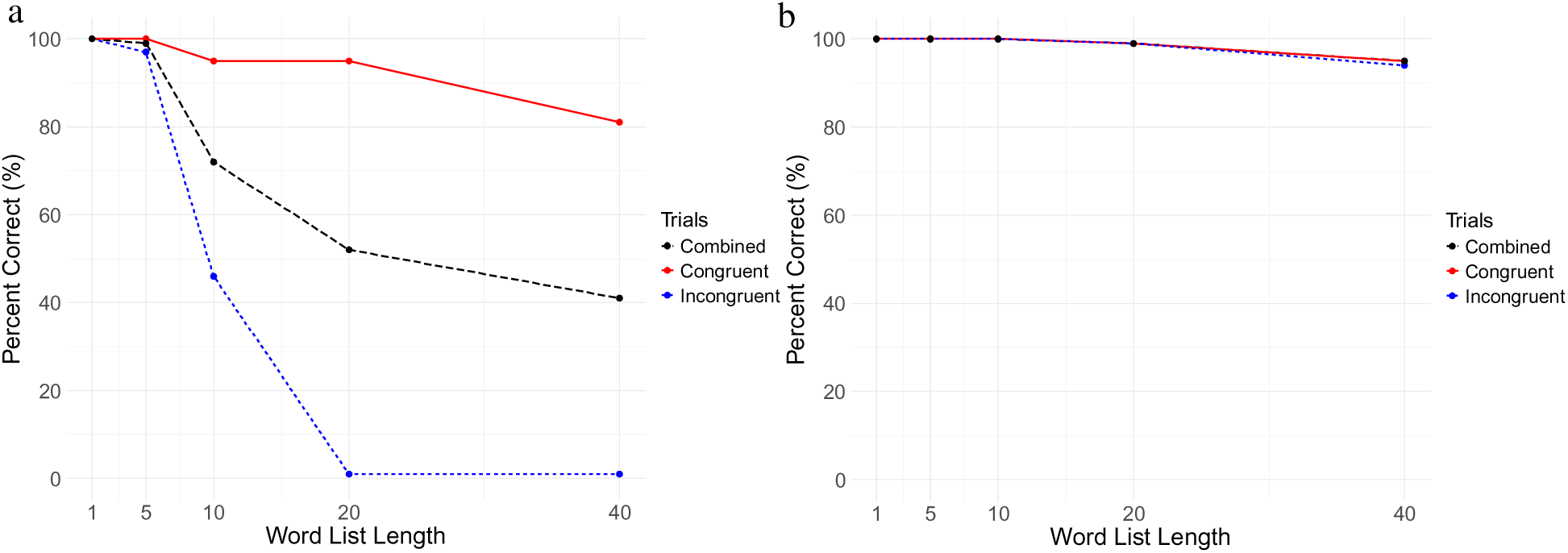
Percent correct for the congruent and incongruent trials in the Mixed condition by word list length (ChatGPT 4o). (a) color naming; (b) word reading.

**Figure 5.**
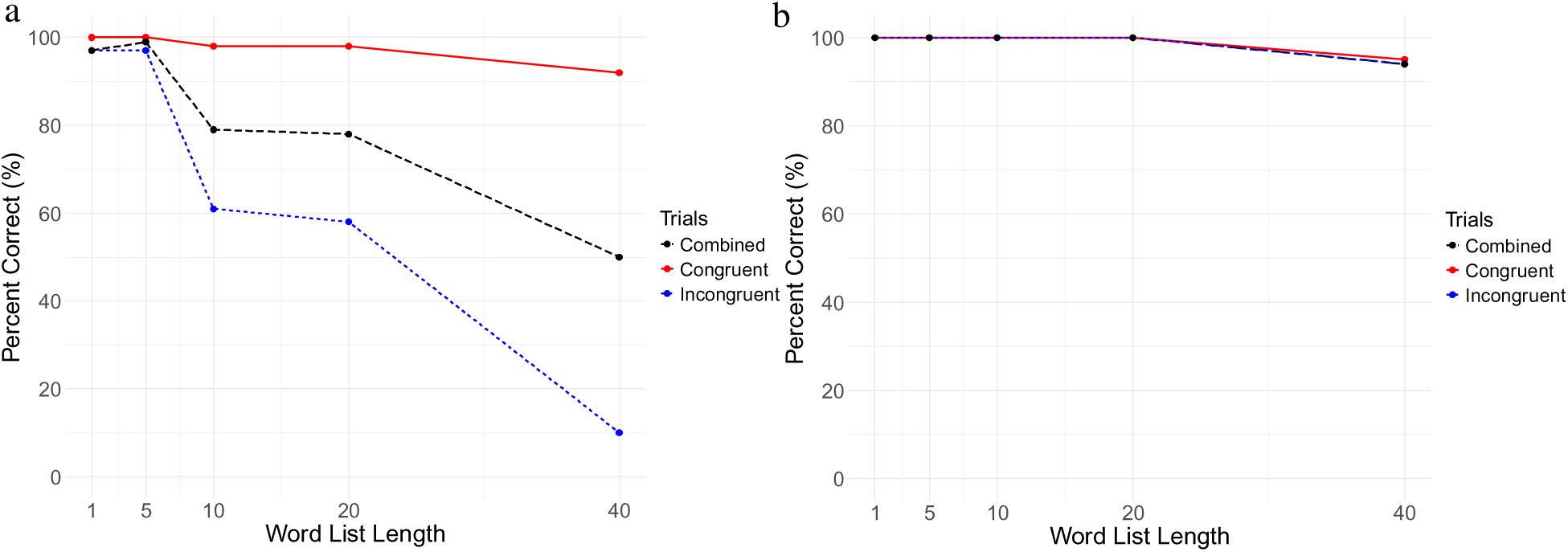
Percent correct for the congruent and incongruent trials in the Mixed condition by word list length (Claude Sonnet 3.5). (a) color naming; (b) word reading.

Claude Sonnet’s 3.5 performance pattern also provides compelling evidence of the conflict effect as a function of list length (Figure 5a). In the mixed trial condition for the overall performance, the model exhibited a progressive decline in accuracy as lists grew longer, starting at a near-perfect 99% accuracy with 5-word lists, dropping to 79% with 10-word lists, and then maintaining relatively robust performance at 78% accuracy with 20-word lists. The model’s performance finally degraded to chance levels (50%), and the responses were only based on word reading but not the color when processing 40-word lists. This gradual degradation pattern, before reaching chance levels, suggests a more robust handling of conflicting information compared to the other model. While Sonnet 3.5 demonstrated better resilience to the incongruent conditions, it still showed impaired performance, maintaining 61% accuracy with 10-word lists and 58% with 20-word lists before declining significantly to 10% accuracy with 40-word lists. Although Sonnet 3.5’s performance degraded at a slower rate, it still exhibited a collapse in executive control that was not seen in neurotypical human trials.

For the neutral condition, GPT4o performed with high accuracy for the list lengths of 1, 5, and 10, with a significant drop for list lengths of 20 (to 75%) and 40 (to 32%) (Figure 2a). The pattern of Claude Sonnet 3.5 is similar (with a notable exception of 73% for a list length of 1): dropped to 78% and 27% for a word list length of 20 and 40, respectively (Figure 3a). The significantly low performance of both LLMs on the longer lists of 20 and 40 items, in contrast to human performance, suggests that these models exhibit a capacity limit in selective attention, which is typically supported by executive control in humans. The predominant error patterns observed on neutral trials manifested in two distinct ways: response omissions, where LLMs completely skipped responding to some of the neutral words, and misattributed color designations, where LLMs incorrectly assigned colors to neutral stimuli (e.g., identifying the word "PEN" as being displayed in blue). While responses occasionally defaulted to neutral words, these instances were rare.

To investigate potential word-specific interference effects, we included a nonword neutral control condition with nonword strings containing only characters of ’X’. The results (Figures 2a&3a) demonstrated a mixed pattern: both GPT-4 and Claude 3.5 Sonnet showed significantly improved performance on color naming tasks with these nonword stimuli at 40 words. GPT-4 exhibited notably better accuracy, while Claude 3.5 Sonnet achieved perfect scores across all the lengths in color naming. Yet for GPT 4o, there was significantly lower performance at 10 and 20 words. These findings suggest that the presence of meaningful words creates interference in the original neutral word condition at longer context lengths.

### Word reading task results

Results for the word reading task demonstrated exceptional performance in both models (99-100% accuracy) across word list lengths from 1 to 20 words (Table 2, Figures 2b&3b), suggesting robust word processing capabilities under various conditions. This is consistent with the pattern of humans. However, at the 40-word list length, performance patterns revealed subtle but distinct model-specific degradations. In congruent trials, both models maintained strong performance with a minimal decline from 20 to 40 words, with 96% for GPT4o and 95% for Sonnet 3.5. Similar modest degradation was observed in mixed trials with 95% for GPT4o and 94% for Sonnet 3.5 at 40 the word list length.

**Table 2.** Results of word reading task (Mean, SD, and SE)

| Condition | Word List Length | Percent Correct - GPT 4o | Percent Correct - Sonnet 3.5 | ChatGPT 4o | Claude Sonnet 3.5 |
| --- | --- | --- | --- | --- | --- |
| Congruent | 1 | 100% | 100% | 1 (0) [0] | 1 (0) [0] |
| Congruent | 5 | 100% | 100% | 5 (0) [0] | 5 (0) [0] |
| Congruent | 10 | 100% | 100% | 10 (0) [0] | 10 (0) [0] |
| Congruent | 20 | 100% | 100% | 19.97 (0.18) [0.03] | 20 (0) [0] |
| Congruent | 40 | 96% | 95% | 38.33 (2.22) [0.4] | 38.1 (1.35) [0.25] |
| Incongruent | 1 | 100% | 100% | 1 (0) [0] | 1 (0) [0] |
| Incongruent | 5 | 100% | 100% | 5 (0) [0] | 5 (0) [0] |
| Incongruent | 10 | 100% | 100% | 10 (0) [0] | 10 (0) [0] |
| Incongruent | 20 | 99% | 100% | 19.73 (0.45) [0.08] | 19.93 (0.37) [0.07] |
| Incongruent | 40 | 92% | 90% | 36.63 (4.97) [0.91] | 35.97 (6.83) [1.25] |
| Mix | 1 | 100% | 100% | 1 (0) [0] | 1 (0) [0] |
| Mix | 5 | 100% | 100% | 5 (0) [0] | 5 (0) [0] |
| Mix | 10 | 100% | 100% | 10 (0) [0] | 10 (0) [0] |
| Mix | 20 | 99% | 100% | 19.8 (0.41) [0.07] | 20 (0) [0] |
| Mix | 40 | 95% | 94% | 37.83 (2.21) [0.4] | 37.73 (1.84) [0.34] |
| Neutral | 1 | 100% | 100% | 1 (0) [0] | 1 (0) [0] |
| Neutral | 5 | 100% | 100% | 5 (0) [0] | 5 (0) [0] |
| Neutral | 10 | 100% | 100% | 10 (0) [0] | 10 (0) [0] |
| Neutral | 20 | 100% | 100% | 19.97 (0.18) [0.03] | 20 (0) [0] |
| Neutral | 40 | 96% | 87% | 38.23 (4.4) [0.8] | 34.63 (4.63) [0.85] |

**Table 3.** Component level summary statistics for the Mix condition of the color naming and word reading tasks.

| Task | Condition | Word List Length | Percent Correct - GPT 4o | Percent Correct - Sonnet 3.5 | ChatGPT 4o | Claude Sonnet 3.5 |
| --- | --- | --- | --- | --- | --- | --- |
| **Color Naming** | Mix | **1** | 100% | 97% | 1 (0) [0] | 0.97 (0.18) [0.03] |
|  | Congruent | 0 - 1 | 100% | 100% | 1 (0) [0] | 1 (0) [0] |
|  | Incongruent | 0 - 1 | 100% | 97% | 1 (0) [0] | 0.97 (0.18) [0.03] |
|  | Mix | **5** | 99% | 99% | 4.93 (0.25) [0.05] | 4.93 (0.25) [0.05] |
|  | Congruent | 2 - 3 | 100% | 100% | 2.5 (0.00) [0.00] | 2.5 (0.00) [0.00] |
|  | Incongruent | 2 - 3 | 97% | 97% | 2.43 (0.00) [0.00] | 2.43 (0.00) [0.00] |
|  | Mix | **10** | 72% | 79% | 7.23 (1.72) [0.31] | 7.93 (1.78) [0.32] |
|  | Congruent | 5 | 95% | 98% | 4.77 (0.00) [0.00] | 4.90 (0.71) [0.13] |
|  | Incongruent | 5 | 46% | 61% | 2.30 (1.70) [0.31] | 3.03 (1.34) [0.25] |
|  | Mix | **20** | 52% | 78% | 10.47 (1.31) [0.24] | 15.63 (2.57) [0.47] |
|  | Congruent | 10 | 95% | 98% | 9.53 (0.65) [0.12] | 9.77 (0.00) [0.00] |
|  | Incongruent | 10 | 1% | 58% | 0.93 (1.11) [0.20] | 5.83 (2.10) [0.38] |
|  | Mix | **40** | 41% | 50% | 16.4 (4.55) [0.83] | 19.9 (4.08) [0.74] |
|  | Congruent | 20 | 81% | 92% | 16.10 (4.36) [0.80] | 18.30 (3.85) [0.70] |
|  | Incongruent | 20 | 1% | 10% | 0.23 (0.57) [0.10] | 2.00 (3.87) [0.71] |
| **Word Reading** | Mix | **1** | 100% | 100% | 1 (0) [0] | 1 (0) [0] |
|  | Congruent | 0 - 1 | 100% | 100% | 1 (0) [0] | 1 (0) [0] |
|  | Incongruent | 0 - 1 | 100% | 100% | 1 (0) [0] | 1 (0) [0] |
|  | Mix | **5** | 100% | 100% | 5 (0) [0] | 5 (0) [0] |
|  | Congruent | 2 - 3 | 100% | 100% | 2.5 (0) [0] | 2.5 (0) [0] |
|  | Incongruent | 2 - 3 | 100% | 100% | 2.5 (0) [0] | 2.5 (0) [0] |
|  | Mix | **10** | 100% | 100% | 10 (0) [0] | 10 (0) [0] |
|  | Congruent | 5 | 100% | 100% | 5 (0) [0] | 5 (0) [0] |
|  | Incongruent | 5 | 100% | 100% | 5 (0) [0] | 5 (0) [0] |
|  | Mix | **20** | 99% | 100% | 19.8 (0.41) [0.07] | 20 (0) [0] |
|  | Congruent | 10 | 99% | 100% | 9.90 (0) [0] | 10.0 (0) [0] |
|  | Incongruent | 10 | 99% | 100% | 9.90 (0) [0] | 10.0 (0) [0] |
|  | Mix | **40** | 95% | 94% | 37.83 (2.21) [0.4] | 37.73 (1.84) [0.34] |
|  | Congruent | 20 | 95% | 95% | 19.89 (0.81) [0.15] | 19.91 (1.06) [0.19] |
|  | Incongruent | 20 | 94% | 94% | 19.88 (0.79) [0.14] | 19.88 (1.05) [0.19] |

For word reading, notable differences emerged in the neutral and incongruent conditions. Claude 3.5 Sonnet exhibited parallel performance declines in both conditions, dropping from 100% at 20 word length to 87% in neutral trials and to 90% in incongruent trials. Conversely, GPT-4o’s performance remained more stable across conditions, with its largest decline occurring in the incongruent condition, from 99% at 20 words to 92% at 40 words. The GPT4o model maintained remarkably consistent performance across other 40-word conditions (congruent: 96%, mixed: 94%, neutral: 96%).

## Discussion

The present study reveals both similarities and differences between human cognitive processes and LLMs’ transformer attention in their response to the Stroop paradigm. The robust appearance of the Stroop effect in the two well-trained state-of-the-art LLMs is noteworthy.

Consistent with classical human studies (MacLeod, 1991; Stroop, 1935), our LLM participants demonstrated reduced performance in the incongruent condition compared to the neutral and congruent conditions - a clear manifestation of the Stroop effect. This performance pattern suggests that the attention mechanisms implemented in transformer architectures exhibit within-modal and cross-modal interference effects analogous to those observed in the human executive control network, likely due to the shared latent space. Moreover, the models’ strong performance on congruent trials for color naming, paired with near-perfect word reading accuracy at equivalent list lengths, indicates a bias toward prioritizing word reading over color naming - likely a result of extensive training in language tasks over other modalities.

However, an important finding is the rapid deterioration of the models’ color naming performance to chance levels as word list lengths increase. This pattern sharply contrasts with human performance, which typically exhibits an interference effect that is insensitive to list lengths. These findings suggest while the LLMs possess exceptional processing capabilities for word reading, even in incongruent conditions, they exhibit significant deficits in color naming for incongruent trials. This highlights a critical limitation in their ability to perform complex tasks requiring conflict resolution and response inhibition.

The near-perfect results in word reading, essentially an optical character recognition (OCR) task, indicate that perception of an extended 40-word list was not hindering performance on the incongruent conditions. The inability to resolve cognitive conflicts in the color naming task due to limited executive control appears to be the primary hindering factor in model performance. Both Claude 3.5 Sonnet and GPT-4 perform significantly below human benchmarks in overall accuracy, with a distinctive pattern of degradation that sees their performance drop below chance levels at 40-word lists, a task length where humans maintain substantially better performance (Dallaway, Lucas, & Ring, 2023; Salihu et al., 2023).

The parallel distributed processing model of the Stroop effect (Cohen, Dunbar, & McClelland, 1990) provides a framework where automaticity strength directly correlates with training exposure duration. This maps onto LLMs since, like humans, LLMs have received substantially more training in word reading than color naming due to their extensive exposure to text. This training asymmetry potentially creates a similar cascading attention mechanism, where the dominant word-reading pathway interferes with the weaker color-naming pathway, ultimately mirroring the automaticity patterns observed in human cognitive processing. This parallel suggests that despite their architectural differences from biological attention networks, LLMs may develop analogous processing hierarchies (Keller et al., 2023) through their training exposure. Additional support comes from the utilization of the backpropagation algorithm (Rumelhart, Hinton, & Williams, 1986), still used in training AI systems, to adjust the strengths of the color naming or word reading pathways.

In humans, research has shown the Stroop interference effect manifests across a clear developmental trajectory. The effect emerges around ages 6-7 years as reading becomes increasingly automatized (Comalli, Wapner, & Werner, 1962). By ages 9-11, children show similar interference reaction times to adults, though with higher error rates (Wright, 2017). The interference effect then remains stable through early and middle adulthood before showing increases after age 60 (Davidson, Zacks, & Williams, 2003; Ward et al., 2021). Research on the relationship between cognitive control (which is a higher-level psychological construct of executive control) and intelligence has demonstrated that cognitive control was more strongly correlated with fluid intelligence than with crystallized intelligence (Chen et al., 2019), a pattern that mirrors the performance characteristics of LLMs. While LLMs excel at tasks requiring crystallized intelligence, such as passing the bar exam to become a lawyer (Katz, Bommarito, Gao, & Arredondo, 2024) and passing medical licensing exams (Bommineni et al., 2023), they still struggle with simple yet novel letter-counting problem-solving tasks (Fu, Ferrando, Conde, Arriaga, & Reviriego, 2024) which are supported by fluid intelligence.

The remarkably poor accuracy in the neutral condition (GPT-4o: 32%; Claude 3.5 Sonnet: 27%) suggests that color naming is a disproportionately demanding cognitive task with a result pattern that is distinctly different from human performance. These neutral word trials, where LLMs correctly process some neutral words while misattributing colors to others, can be characterized as a form of hallucination (Huang et al., 2023), a phenomenon where LLMs have a tendency to generate unfounded associations. Furthermore, the extended sequences of unrelated words appeared to trigger confabulatory interference effects (Sui, Duede, Wu, & So, 2024; Smith, Greaves, & Panch, 2023), where the model’s ability to maintain task-relevant goals becomes compromised. Similar to human executive control mechanisms, LLMs demonstrate impaired interference suppression during longer sequences, suggesting analogous limitations in maintaining task-relevant attention over extended contexts (Kane & Engle, 2003).

The cognitive impairment exhibited by LLMs, though, likely stems from the architecture of transformer models, particularly their embedding space topology. Unlike human cognitive systems, where the semantic relationship between neutral words and color terms has minimal impact on color perception tasks, transformer models encode these relationships in their high-dimensional latent space geometry. Specifically, the soft attention mechanism assigns a probability to the entire context and incorporates the irrelevant information from the neutral words’ semantic context into its latent representation (Welleck et al., 2019; Weston & Sukhbaatar, 2023). This degradation in accuracy for next token generation is a transformer-specific interference effect that produces low accuracy when models attempt to perform color identification on extended sequences of semantically unrelated words. In comparison, interference in human Stroop effect experiments is reduced by inducing great effort for dynamic modulation and response inhibition (Cohen, Dunbar, & McClelland, 1990) or modulation of input by blurring the stimuli (Pálfi, Parris, Collins, & Dienes, 2019) or even via hypnotic suggestions (Parris, Hasshim, & Dienes, 2021; Raz, Shapiro, Fan, & Posner, 2002). The embedding space architecture’s potential performance limitations mirror aspects of human cognition, where semantic processing is an uncontrollable endogenous form of attention (Kinoshita et al., 2018), yet can be modulated through executive control mechanisms to filter distracting word meaning. The broader construction of cognitive control suggests that executive control is part of an integrated system in which attention networks work together to process information and reduce uncertainty, rather than being a standalone control mechanism (Fan, 2014).

In our exploratory trials without explicit prompts, Claude 3.5 Sonnet demonstrated recognition of the Stroop paradigm but still produced incorrect responses despite output with word-color relationship mappings (see Supplementary Figure 1). This behavior has been investigated with experiments where the models recognize ten classic illusions (Ullman, 2024), further providing evidence that the stimuli alone can trigger task recognition and orientation in these AI systems, even absent explicit prompts. However, the persistence of incorrect responses despite task recognition suggests a dissociation between task understanding and execution capabilities. The models’ effective context window for executive control is notably shorter than their general processing capacity, with performance degrading in conflict conditions at just 10 words - a fundamental limitation that must be addressed to achieve parity with biological attention systems.

The ultimate goal of AI research is to develop artificial general intelligence (Bubeck et al., 2023) comparable to human abilities, and human cognitive development provides crucial insights for this endeavor. While what we traditionally observed as executive control may be a fundamental manifestation of general intelligence rather than a distinct cognitive domain (Royall & Palmer, 2014), research has also shown that early attentional control abilities in young children establish the foundation for later-emerging of a higher-level construct called executive functions (Miyake et al., 2000; Garon, Bryson, & Smith, 2008). Human cognitive development begins with basic attention control in childhood, which then scaffolds three core executive functions: inhibitory control, working memory, and cognitive flexibility, ultimately enabling complex abilities like problem-solving and self-monitoring (Diamond, 2013). This developmental trajectory suggests that AI systems, like humans, may need to master fundamental attention mechanisms, e.g., executive control, before achieving the complex reasoning and generalized problem-solving capabilities characteristic of mature executive functions.

A recent study examined LLMs’ cognitive capabilities using the Montreal Cognitive Assessment, which includes the Stroop effect, revealed mild cognitive impairment patterns in both GPT-4 and Claude 3.5 Sonnet, extending previous findings regarding specific cognitive limitations in AI systems, particularly in domains of emotional processing and empathy (Patel & Fan, 2023). The patterns suggest homologous mechanisms, specifically regarding the established correlation between alexithymia and executive functions throughout human development.

Similar to biological attention and control systems, where attention serves as a fundamental mechanism for flexibly controlling limited computational resources and integrating multiple cognitive processes, including awareness, vigilance, saliency, and executive control (Lindsay, 2020), transformer attention systems may also implicitly develop functionally similar hierarchical networks (Keller et al., 2023) that support executive functions and general intelligence (Barbey et al., 2012). These parallels offer important insights into machine consciousness research (e.g., Dehaene, Lau, & Kouider, 2017).

Recent transformer architecture innovations, such as those described in a Google Research paper (Behrouz, Zhong, & Mirrokni, 2024), focus primarily on enhancing memory capabilities. However, this approach may not address the core limitations in attention mechanisms, specifically, the need for sophisticated alerting, orienting, and executive control networks that enable cognitive flexibility. Human cognition, for instance, employs a task set to manage extended contexts despite limited working memory, maintaining consistent performance in Stroop tasks regardless of list length. While these innovations in transformer architectures can dramatically improve context window sizes, they may be missing crucial elements of biological attention systems. The key limitation appears not to be memory capacity but rather the absence of robust attention mechanisms that enable effective goal-directed behavior. Future developments might benefit from implementing more sophisticated executive control systems that can handle decision conflicts through structured, goal-directed processing rather than relying solely on enhanced memory capabilities.

