## Supplementary Materials for "Deficient Executive Control in Transformer Attention"

Supplementary Table 1. Task duration and list length effects on reaction time (ms) and accuracy (%) on the color naming task.

| Task Duration | Word List Length | Condition | Correct (%) | Reaction Time (ms) |
|---|---|---|---|---|
| 5-min[1] | 120 | Congruent | 97.20 ± 1.83 | 594 ± 84 |
| | 120 | Incongruent | 95.20 ± 3.00 | 885 ± 144 |
| 10-min[1] | 240 | Congruent | 95.43 ± 2.81 | 595 ± 75 |
| | 240 | Incongruent | 95.87 ± 2.65 | 834 ± 145 |
| 20-min[1] | 480 | Congruent | 96.23 ± 3.18 | 580 ± 95 |
| | 49 | Incongruent | 95.92 ± 2.89 | 817 ± 122 |
| 60-min[2] | | | | |
| *Scoring Block 1* | 500 | | 97.30 ± .93 | 900 ± 50 |
| | 125 | Congruent | N/A | N/A |
| | 375 | Incongruent | N/A | N/A |
| *Scoring Block 2* | 500 | | 97.2 ± 1.06 | 920 ± 45 |
| | 125 | Congruent | N/A | N/A |
| | 375 | Incongruent | N/A | N/A |
| *Scoring Block 3* | 500 | | 97.30 ± .96 | 940 ± 50 |
| | 125 | Congruent | N/A | N/A |
| | 375 | Incongruent | N/A | N/A |

Note. Adapted from “Investigating the influence of prolonged Stroop task on mental fatigue and the consequences on corticospinal and corticocortical excitability: A pilot study,” by A. T. Salihu, K. D. Hill, M. Zoghi, and S. Jaberzadeh, 2023, medRxiv. https://doi.org/10.1101/2023.11.28.23299165. CC BY-NC.

"Effects of Stroop task duration on subsequent cognitive and physical performance," by N. Dallaway, S. J. Lucas, and C. Ring, 2023, Psychology of Sport and Exercise, 68, 102459. Copyright 2023 by Elsevier.

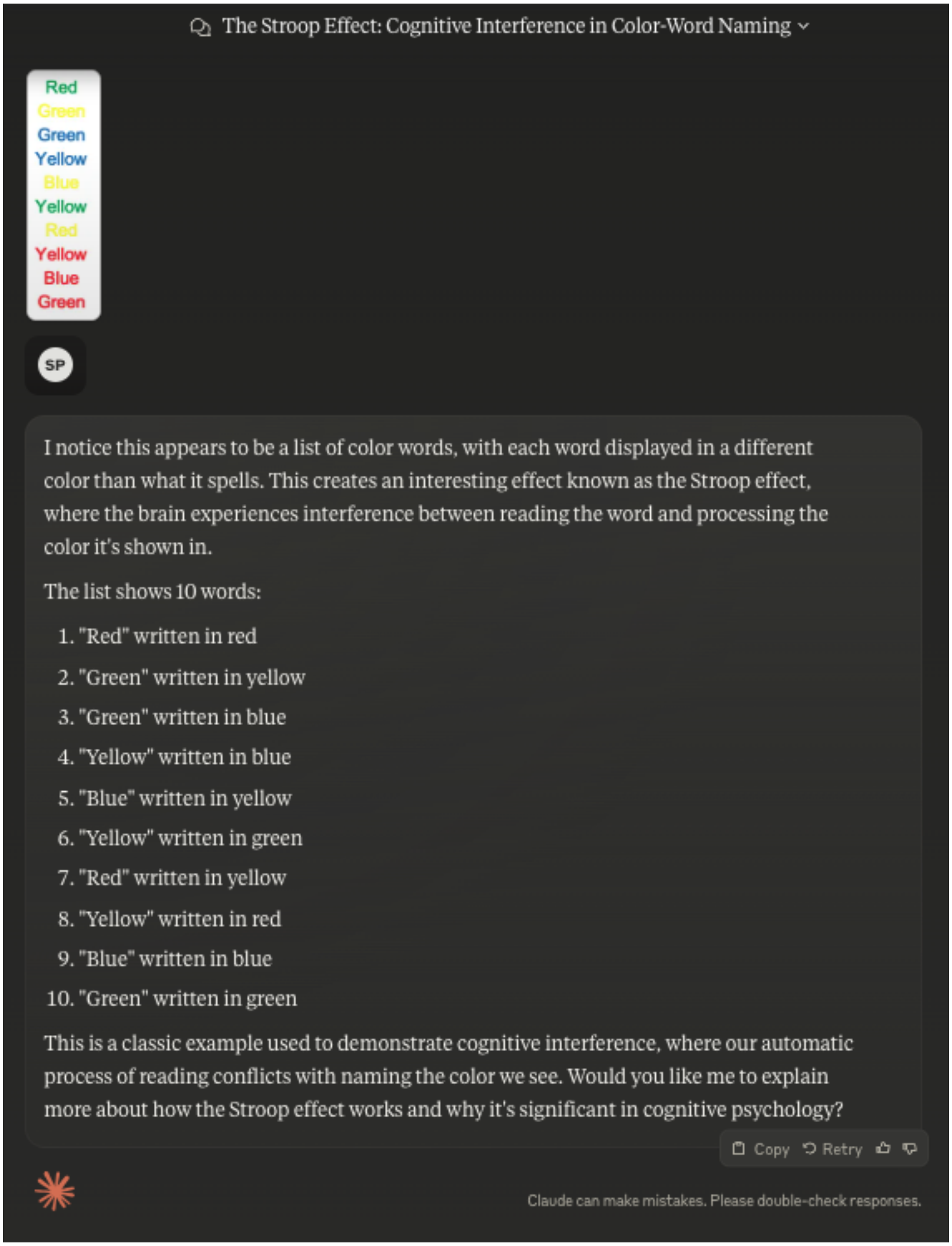


Figure 1. Claude Sonnet 3.5 recognizes the Stroop effect without an explicit prompt yet still provides only 80% accuracy
